# CreLite: An Optogenetically Controlled Cre/loxP System Using Red Light

**DOI:** 10.1101/823971

**Authors:** Shuo-Ting Yen, Kenneth A. Trimmer, Nader Aboul, Rachel D. Mullen, James C. Culver, Mary E. Dickinson, Richard R. Behringer, George T. Eisenhoffer

## Abstract

Precise manipulation of gene expression with temporal and spatial control is essential for functional studies and the determination of cell lineage relationships in complex biological systems. The Cre-*lox*P system is commonly used for gene manipulation at desired times and places. However, specificity is dependent on the availability of tissue- or cell-specific regulatory elements used in combination with Cre or CreER (tamoxifen-inducible). Here we present CreLite, an optogenetically-controlled Cre system using red light in developing zebrafish embryos. Cre activity is disabled by splitting Cre and fusing the inactive halves with the *Arabidopsis thaliana* red light-inducible binding partners, PhyB and PIF6. In addition, PhyB-PIF6 binding requires phycocyanobilin (PCB), providing an additional layer of control. Upon exposure to red light (660 nm) illumination, the PhyB-CreC and PIF6-CreN fusion proteins come together in the presence of PCB to restore Cre activity. Red-light exposure of transgenic zebrafish embryos harboring a Cre-dependent multi-color fluorescent protein reporter (*ubi:zebrabow*) injected with CreLite mRNAs and PCB, resulted in Cre activity as measured by the generation of multi-spectral cell labeling in various tissues, including heart, skeletal muscle and epithelium. We show that CreLite can be used for gene manipulations in whole embryos or small groups of cells at different stages of development. CreLite provides a novel optogenetic tool for precise temporal and spatial control of gene expression in zebrafish embryos that may also be useful in cell culture, *ex vivo* organ culture, and other animal models for developmental biology studies.

## INTRODUCTION

The study of gene function and cellular lineage relationships in rapidly changing environments such as the developing embryo, regeneration of new tissue after injury and changes during carcinogenesis requires accurate spatial and temporal control of genome manipulation. Cyclic recombinase (Cre) originates from bacteriophage P1 and has been applied to specifically target defined *lox*P sites to manipulate the genome at a specific time and place (Feil et al., 1997; Lee et al., 1999). The versatility of the Cre/*lox*P system has been broadly applied for conditional control of genome manipulation, yet specificity is currently dependent on the availability of tissue- or cell-specific regulatory elements used in combination with Cre or drug-inducible Cre systems. Cell or tissue types without known regulatory elements at desired stage of development or regeneration, or those sensitive to drug treatment, will be difficult to study.

Optogenetic approaches have recently emerged as a new methodology that allows both temporal and spatial control of biological processes in a precise manner. Phytochrome-like proteins and their binding partners found in the plant *Arabidopsis thaliana* and bacterium *Rhodopseudomonos palustris* have provided a wide spectrum of light-inducible protein binding pairs for possible optogenetic use, including blue light (Cry2 and CIB), red light (PhyB and PIF), and near-infrared light (BphP1 and PpsR2) (Kaberniuk et al., 2016; Kennedy et al., 2010a; Levskaya et al., 2009). Among these light-inducible binding pairs, blue-light based systems have wide applications in cell culture or superficial tissues. However, the red-light induced PhyB and PIF, is preferable as a light sensor for optogenetic manipulation in light-sensitive embryonic tissue or deep tissues due to the low photo-toxicity and great tissue penetrance of red light. As a result, we designed a new red-light inducible Cre/*lox*P system, we call the CreLite system, to provide precise spatial and temporal control of genetic manipulation in living tissues.

The CreLite system consists of two parts, the light sensing module and the recombinase module. The light-sensing module is the red-light inducible binding pair, PhyB and PIF6. In *Arabidopsis*, PhyB binds to PIF6 in the presence of its prosthetic group, phytochromobilin (PΦB), in response to red light (~650 nm). These proteins then act as co-transcriptional activators of target genes. PhyB and PIF family members have been successfully engineered to create light-controlled gene expression systems (Beyer et al., 2015; Buckley et al., 2016; Levskaya et al., 2009; Müller et al., 2013a; Noda and Ozawa, 2018). In the recombinase module, Cre is split into N- and C-terminal regions that separately lack Cre activity. However, when these regions are physically brought together, Cre activity is restored (Jullien et al., 2003). Therefore, we created fusion proteins of PhyB and PIF6 with the split Cre complementation system, PhyBCreC and PFI6CreN. Notably, the requirement of prosthetic group PΦB in this system provides more control of this system and avoids keeping animals continuously in the dark. Practically, PΦB is usually substituted by its analog phycocyanbilin (PCB) due to its broad availability. In the absence of PCB and red light, PhyBCreC and PIF6CreN will not bind each other and thus there will be no Cre activity. However, in the presence of PCB and red light, PhyBCreC and PIF6CreN will bind each other, leading to Cre activity.

We tested the function of the CreLite system in zebrafish. Red-light exposure of transgenic zebrafish embryos harboring a multi-color fluorescent Cre reporter (*ubi:zebrabow*) and injected with *CreLite* mRNAs and PCB resulted in Cre activity and the generation of multi-spectral cell labeling in various tissues, including eyes, skeletal muscle and epithelium. Our results also show that CreLite can be used for activation of whole-embryos or small groups of cells at different times throughout development. All the associated plasmids for CreLite have been deposited in publically available databases and provide new tools for precise temporal and spatial control of gene expression in cell culture, *ex vivo* organ culture, and animal models for developmental biology studies.

## RESULTS AND DISCUSSION

### Design and functional test of the light inducible Cre (CreLite) system in zebrafish embryos

To create a light inducible Cre system, we split Cre into two fragments (N-terminus Cre and C terminus Cre) and fused them with the red light inducible dimer, PhyB and PIF6 (Figure 1.). In this system, the split Cre should not show activity until exposed to red-light (around 650 nm) with PCB. Specifically, the synthetic fusion proteins consist of the photo-sensory module of phytochrome B (1-621 aa; NTE-PAS-GAF-PHY) (Buckley et al., 2016) and the C-terminus Cre (60-343 aa), and the active protein binding (ABP) domain of PIF6 (1-100 aa) and N-terminus Cre (19-59 aa) (Jullien et al., 2003), referred to as PhyBΔCreC and PIF6CreN (Figure 1). Both of the two fusion proteins have an N-terminal SV40 nuclear localization sequence (NLS) to ensure nuclear translocation. The term “CreLite” is used to represent PhyBΔCreC and PIF6CreN together.

**Figure 1.**
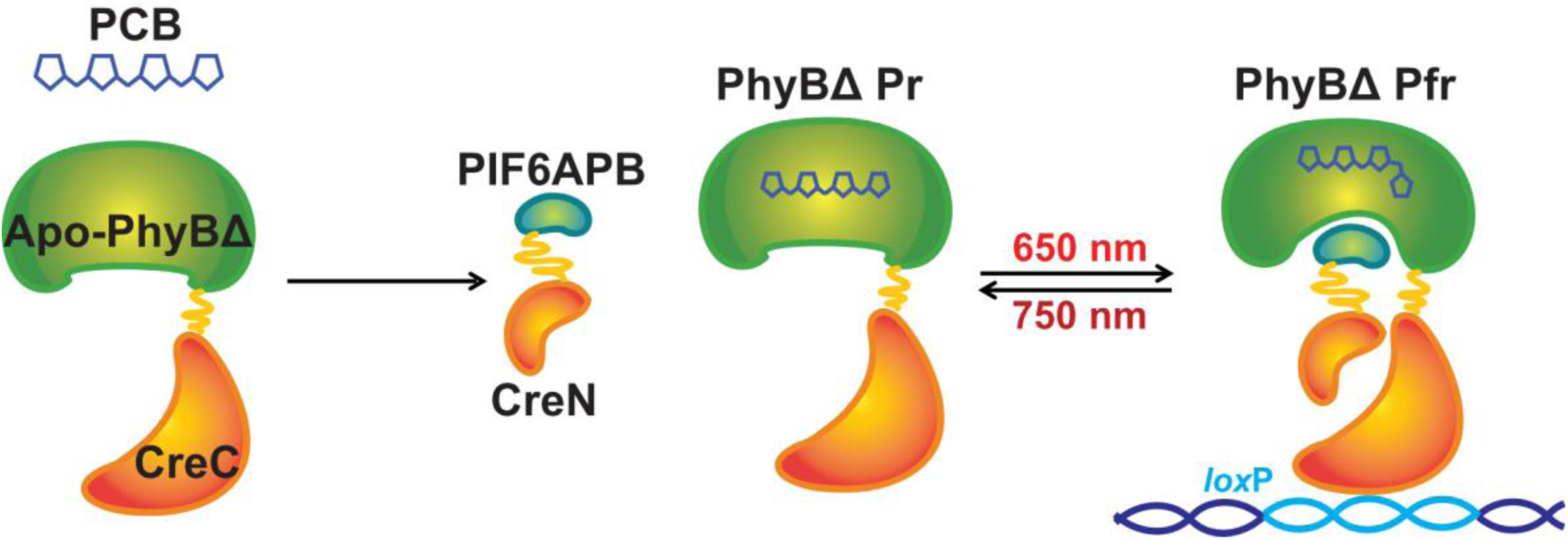
The CreLite system. The light inducible protein binding pair, truncated Phytochrome B (PhyBΔ) and the active protein binding domain of Phytochrome B interacting factor 6 (PIF6APB) are fused with C-terminus Cre (CreC) and N-terminus Cre (CreN), respectively. A cofactor analog, phycocyanobilin (PCB), is required for PhyBΔ to be functional. Once exposed to red light (λ_max_ = 650 nm), the PhyBΔ and PIF6APB bind each other and bring the two halves of Cre together, resuming its recombinase activity.

To test the function of CreLite in zebrafish, we injected *in vitro* transcribed 300 ng/μL *PhyBΔCreC* and 200 ng/μL *PIF6APBCreN* mRNAs along with 1.4 µM PCB into zebrafish embryos *Tg(ubi:zebrabow-S)*^*a132*^ (Pan et al., 2013), referred as *ubi:zebrabow*, at the one cell stage (Figure 2A-B). The *ubi:zebrabow* zebrafish ubiquitously expresses red fluorscent protein (RFP) but upon Cre-mediated recombination can express yellow fluorescent protein (YFP) or cyan fluorescent protein (CFP). Red-light exposure at ~6 hpf (hours post fertilization) of *ubi:zebrabow* injected with CreLite mRNAs and PCB resulted in Cre activity and generated a diverse color profiles in various tissues, including eyes, skeletal muscle and epithelium. We observed 56% of larvae harbored cells with fluorescence conversion from RFP to CFP or YFP (*n* = 89). In contrast, injection mixtures lacking either the *CreLite* mRNAs, PCB, or red light exposure showed little to no fluorescence conversion in 48 hpf embryos (Figure 2D). The resulting Cre reporter expression was found in both deep and superficial tissues, including the epithelia, eyes, muscle, or even the blood vessels (Figure 2E), suggesting penetrance of the red-light deep into the body of the animal and thus the Cre-Lite system could be used for conversion of diverse cell types. Further, we observed very little phototoxicity. The abnormality rate of red light exposed embryos (9.32%; *n* = 195) is not significantly different from the abnormality rate of non-exposed embryos (9.73%; *n* = 129) (Supplementary Figure 1D). The low conversion rate (20%) in the non-exposed embryos is likely due to exposure of ambient light or stochastic dimerization when high concentrations of CreLite proteins exist in cells.

**Figure 2.**
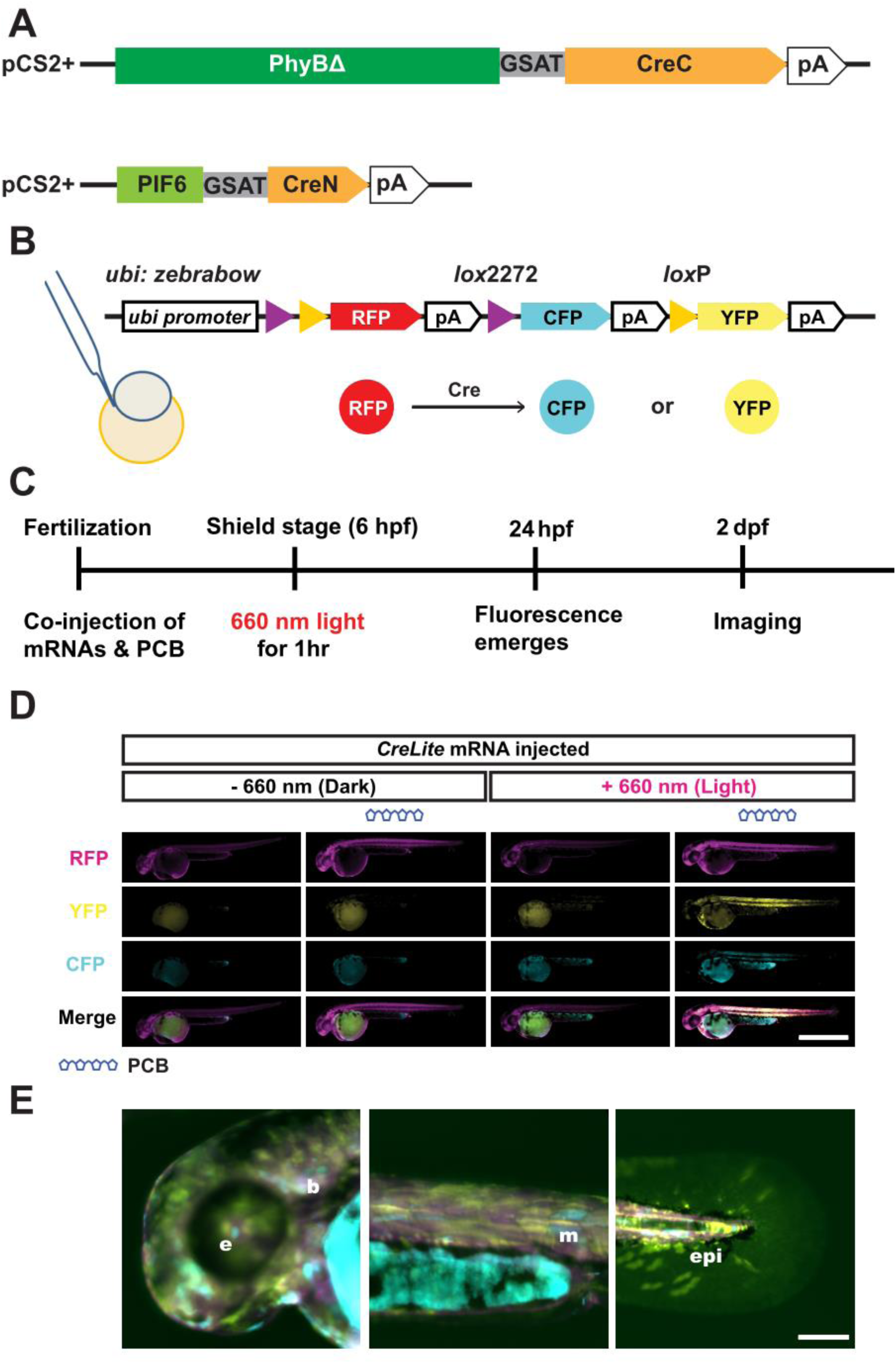
CreLite-mediated recombination in zebrafish embryos. *PhyBΔCreC* and *PIF6CreN* gene were subcloned into pCS2+ expression vector for in vitro transcription (A). The resulting CreLite mRNAs, along with the cofactor PCB, were microinjected into *ubi:zebrabow* zebrafish reporter line (B), which changes its color from red to either cyan and yellow depending on the Cre-mediated recombination event. Injected embryos were exposed to 660 nm red light for an hour at shield stage (C). Negative controls including no PCB and no light exposure were examined (D). Embryos with both PCB and red light exposure show broad recombination in the animal; Scale bar = 1 mm. YFP signal of converted animals shows color conversion in various tissues (E), including brain (b), eyes (e), muscle (m), and epithelial (epi) cells; Scale bar = 100 µm.

### Induced reporter gene expression at different developmental stages and at specific regions of interest

The CreLite system is designed to allow precise spatial and temporal control of gene expression, and therefore, we first examined induction of Cre activity at different time points after injection and exposure to red light. The *ubi:zebrabow* embryos were injected with CreLite mRNAs with PCB were exposed to red light for an hour at different stages of development, 6 hpf, 1 day post fertilization (dpf), and 2 dpf, and the fluorescence conversion (from RFP to CFP or YFP) was assessed at 1 day post exposure, (Figure 3 A-D). Exposure to red light at both 6 hpf and 1 dpf showed a high percent of embryos with fluorescence conversion of distinct cells in individual embryos (~50%). The percent of embryos with fluorescence conversion of distinct cells in individual embryos is reduced (33%) for exposure at 2 dpf. This may be due to the gradual depletion of the injected mRNA as the embryos developed.

**Figure 3.**
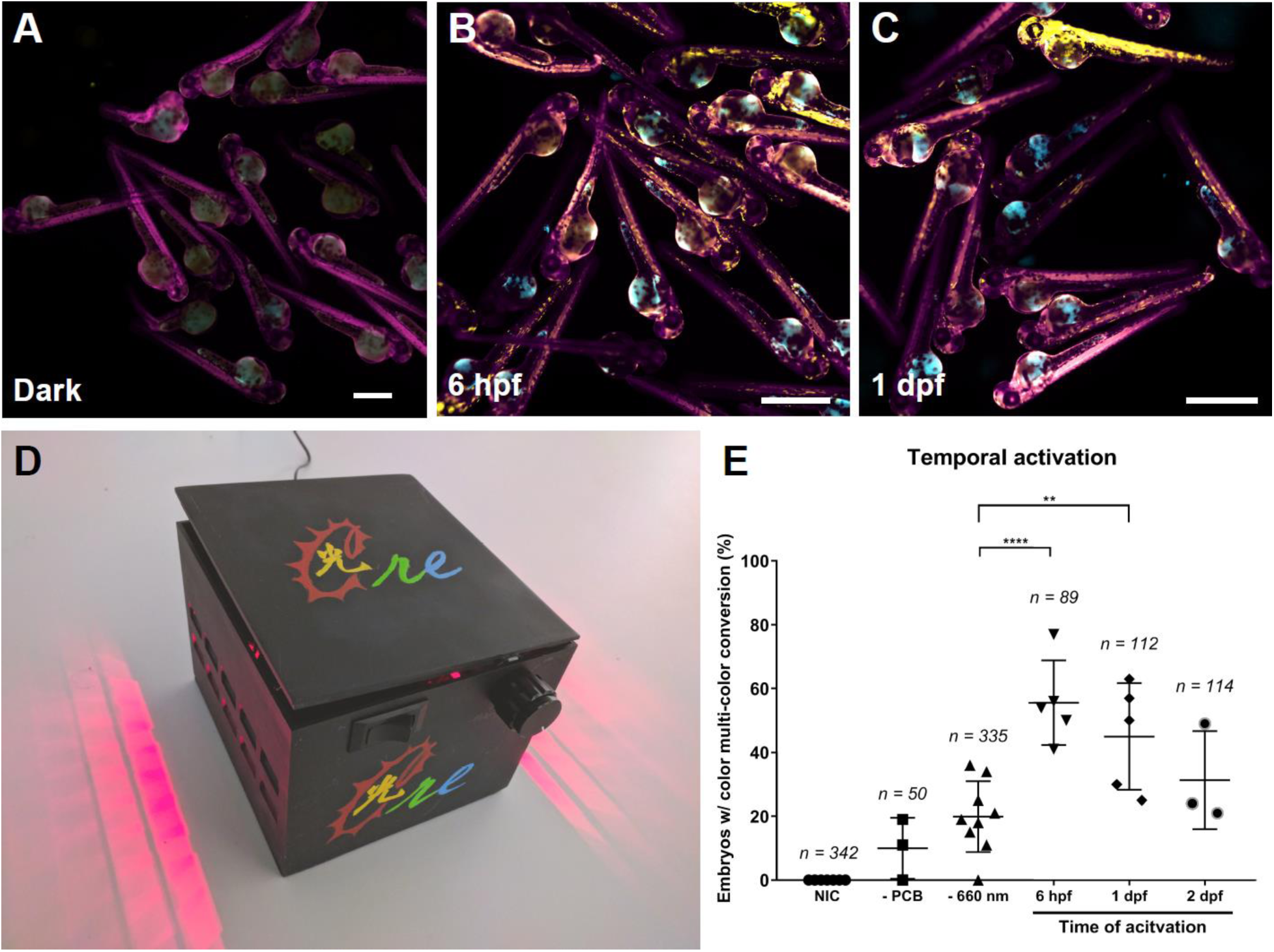
CreLite-mediated recombination at different times in development. *CreLite* mRNAs injected *ubi:zebrabow* embryos were exposed to 660 nm red light in the LED lightbox (D) at different time points (i.e. 6 hpf, 1 dpf) (B and C). Mean number of embryos containing cells that exhibit YFP and/or CFP fluorescence a day after red light exposure were counted (E). Data are from three independent experiments and error bars represent *SD*; **** *p* < 0.0001, ** *p* < 0.01. Ordinary one-way analysis of variance (ANOVA) with Dunnett’s multiple comparisons test. *n* = total number of embryos in each group. Each data point represents a clutch of embryos. NIC: non-injected control. Scale bar = 1 mm.

To test if the CreLite system can be controlled and activated at specific regions of interest in zebrafish, we performed activation on the developing eye region. A 50 µm diameter circle covering a portion of the eye was exposed to red light (Figure 4A-B). Quantification of YFP/CFP signals at 6 hours post exposure (hpe) within the activated region suggests Cre activity was induced successfully at the region of interest, with little impact on surrounding tissues (Figure 4C). We found successful regional activation of the Cre reporter in 12% (*n* = 17 from 2 independent experiments) of all the exposed embryos. These results suggest that regional activation is mosaic and may be due to the distribution of the key components (PhyBΔCreC mRNA, PIF6CreN mRNA, and PCB) not being homogenous in the embryos and thus the co-existence of these components in the same cell may not always be achieved, leading to a mosaic activation after light exposure. In a recent study, a comparable mosaic activation has also been shown (Reade et al., 2017). As a result, we suggest precise temporal and spatial control using CreLite is achievable. In the future, generating a transgenic line carrying CreLite fusion protein genes will help to achieve more even distribution of the CreLite components in embryos and further improve the activation rate after light exposure.

**Figure 4.**
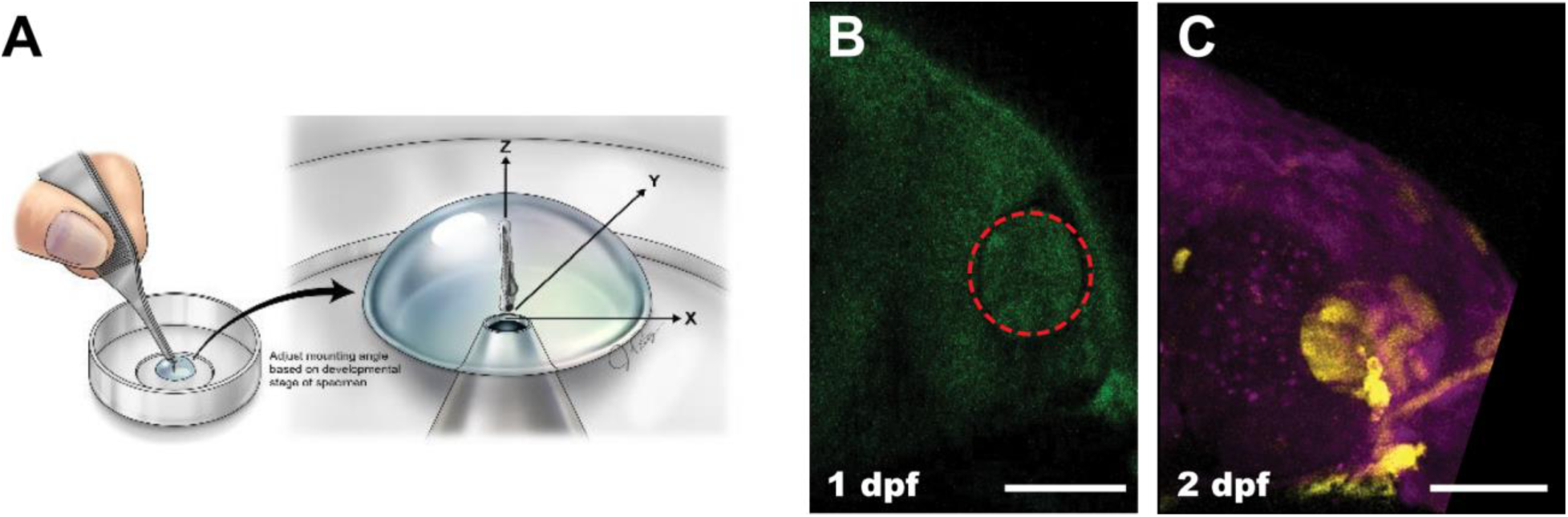
Spatial and temporal recombination via CreLite. Regional locations of the developing eye were activated by exposure to red-light at 1 dpf. CreLite mRNA and PCB injected *ubi:zebrabow* embryos were mounted rostrally (A) in LMP agarose. Regions of interest (red circle in B) were exposed to 640 nm laser. The color conversion was examined at 2 dpf (C); Scale bar = 50 µm.

### Conclusions

Precise spatial and temporal manipulation of gene expression is key for studies of development and tissue regeneration in a wide variety of organisms. Classical methods such as using beads soaked with proteins such as signaling ligands like fibroblast growth factors or small molecules like retinoic acid allows functional studies (Crossley et al., 1996). Yet, these methods are also limited to the studies of secreted molecules and physically placing the soaked beads in the tissue may introduce mechanical disruption or even damage the tissue, leading to undesired side effects. Optogenetic tools have facilitated better control of the expression of their gene of interest, including CRY-CIBN based PA-Cre and LACE systems (Polstein and Gersbach, 2015; Schindler et al., 2015; Taslimi et al., 2016), LOV based TAEL system (Chen et al., 2013; Motta-Mena et al., 2014; Reade et al., 2017; Wang et al., 2012), FKF/GI based system (Quejada et al., 2017) and PhyB-PIF based system (Beyer et al., 2015; Müller et al., 2014, 2013a; Noda andOzawa, 2018). Most of these tools achieve light-inducible gene expression by using viral trans-activator VP16 or VP64, which requires continuous light activation when prolonged gene expression is desired. Here, we successfully demonstrate a red-light inducible Cre/*lox*P system that allows light inducible genome manipulation and constant gene expression after induction. In combination with *lox*P alleles (e.g. *lox*P-stop-*lox*P, *brainbow*, *MADM* or *FLEX*), it is also possible to achieve switching of multiple genes with precise regional light induction for clonal analysis, lineage tracing, or to interrogate carcinogenesis using our system.

Notably, recent studies have demonstrated a CRY-CIBN based blue-light inducible Cre/*lox*P system in cell culture and *Drosophila melanogaster* (Boulina et al., 2013; Kennedy et al., 2010b; Schindler et al., 2015; Taslimi et al., 2016). However, a red-light inducible system has several strengths for geneticists and developmental biologists. The red-light inducible system is intrinsically low photo-toxicity with a high tissue penetrance, as we found YFP or CFP signals in deep tissues (e.g. muscle and heart; Figure 2E) with very few abnormal embryos after prolonged red light exposure (1 hour at 64 µW/mm^2^). The red-light induced recombination event is irreversible, and therefore, continuous light-induction is not required in our system. Moreover, a red-light inducible system frees up more compatible imaging excitation wavelengths (Tischer and Weiner, 2014) and allows blue light excitation for widely used GFP-tagged proteins, which is critical for live-imaging in tissues like developing organs or regenerating tissues.

The co-factor PCB is required for the CreLite system to function. The delivery of PCB into adult or deep tissues may be a concern, and the additional component may not be ideal for cultured cells or other systems that require minimum manipulation with light. However, the latter issue can be addressed by introducing genes encoding enzymes for PCB synthesis (HO1 and PcyA) in the system, which has been shown previously in a cell culture system (Müller et al., 2013b). For *ex vivo* or *in vivo* systems like developing zebrafish embryos, the cofactor as an additional component in this system is actually beneficial and allows an additional level of control, to prevent handling the entire experiment in the dark or under green safe light.

In conclusion, we demonstrate that the CreLite system is a promising red-light inducible Cre/*lox*P system. Our red-light inducible Cre/*lox*P system will be useful for control of gene expression in deep tissues and for live-imaging after Cre-mediated induction. This system also opens up the possibilities for more sophisticated genome manipulation that utilizes both red and blue light to dissect complex biological questions. CreLite provides a novel optogenetic tool for precise temporal and spatial control of gene expression in zebrafish embryos that may also be useful in cell culture, *ex vivo* organ culture, and other animal models for developmental biology studies (The plasmids listed in Table S2 have been generated to facilitate application of the Cre system to other models).

## MATERIAL AND METHODS

### Zebrafish

The zebrafish animal protocol was approved by the Animal Care and Use Committee of the University of Texas MD Anderson Cancer Center. The *Tg(ubi:Zebrabow-S)*^*a132*^ (Pan et al., 2013), referred as *ubi:zebrabow* zebrafish were maintained as both homozygotes and hemizygotes in a 28.5°C fish room with a circulating water system and a regular light/dark cycle of 14 hours light and 10 hours darkness.

### Plasmids

The PIF6CreN fragment containing coding sequence (see supplementary information) was synthesized by GeneArt™ Strings™ DNA fragments service (Thermo Fisher Scientific Inc.) and cloned into pSC-B by StrataClone™ Blunt PCR Cloning Kit (Agilent Technologies, La Jolla, CA). To generate pUBC-PIF6iCreN transgene vector, the PIF6CreN fragment was subcloned into cHS2x-UBC-SH-RG-cHS2x (Stewart et al., 2009) at restriction enzyme sites *Apa*I and *Xho*I. The PIF6CreN fragment was also amplified by primers SY118 and SY119 (Table 1.) and cloned into expression vector pCS2+ at *Eco*RI and *Xho*I sites for *in vitro* transcription.

The pCAGEN-PhyBCreC transgene vector was assembled by a four-way ligation. This ligation reaction combines NLS-PhyB (1-908 aa), GSAT linker, and CreC (60-343 aa) into a mammalian expression vector pCAGEN, which was a gift from Connie Cepko (Addgene plasmid # 11160; http://n2t.net/addgene:11160; RRID:Addgene_11160) (Matsuda and Cepko, 2004). PhyB fragment was PCR amplified from pBS-PhyBCre (a gift from Nader Aboul and Dr. Rachel D. Mullen) with primers SY096 and SY097. CreC fragment was PCR amplified from pOG321 (Addgene plasmid # 17736) (O’Gorman et al., 1997) by primers SY094 and SY095. The middle component containing GSAT linker sequence (Part:BBa_K404301, iGEM Registry of Standard Biological parts) between PhyB and CreC was synthesized by GeneArt™ Strings™ DNA fragments service (Thermo Fisher Scientific Inc.) and cloned into pSC-B by StrataClone™ Blunt PCR Cloning Kit (Agilent Technologies, La Jolla, CA). *Eco*RV, *Sph*I, *Sex*AI, and *Not*I (from 5’ to 3’) restriction sites were used to link all three fragments together into pCAGEN. The resulting PhyBCreC fragments were also subcloned into pCS2+ for *in vitro* transcription. We initially constructed the fusion protein with the long form PhyB (1-908 aa; NTE-PAS-GAF-PHY-PAS-PAS) instead of PhyBΔ (1-621 aa; NTE-PAS-GAF-PHY) in our system. However, a previous study has shown limited fusion protein expression in zebrafish when the long form PhyB (which includes two additional PAS domains and plant nuclear localization sequence) is used (Buckley et al., 2016). As a result, we modified our construct and replaced PhyBCreC by PhyBΔCreC in our following experiments. The pCS2+ PhyBΔCreC construct was made by replacing middle part of the sequence of PhyBCreC (between *Nco*I and *Nru*I) by a 230 bp fragment (Supplementary data) synthesized by GENEWIZ, resulting the short form of PhyB, PhyBΔ (1-621 aa), without PAS-A and PAS-B domains (Buckley et al., 2016).

### Zebrafish embryo microinjection

PhyBΔCreC and PIF6CreN mRNAs were *in vitro* transcribed from pCS2+PIF6CreN (Addgene #131780) and pCS2+ PhyBΔCreC (Addgene #131781) by mMESSAGE mMACHINE^®^ SP6 transcription kit (Ambion). The capped mRNA were purified by alcohol precipitation and resuspended in DEPC-treated water. The microinjection of zebrafish embryos performed as described previously (Eisenhoffer and Rosenblatt, 2011; Yuan and Sun, 2009). Briefly, borosilicate glass capillaries with filaments (OD 1.2mm, ID 0.94, 10cm length; ITEM#: BF120-94-10) were pulled using a Sutter P-97 Pipette Puller to generate the injection needles. The *ubi:zebrabow* (Pan et al., 2013) zebrafish embryos were collected in E3 embryo medium and placed into a 10 cm dish with 3% agarose with troughs. The troughs were made by the microinjection mold (Adaptive Science Tools, # TU-1). Under a Leica S6D LED dissecting stereomicroscope, the mRNAs and 1.4 µM of phycocyanobilin (PCB) (SiChem GmbH, Bremen, Germany) in the KCl injection buffer were co-injected into *ubi:zebrabow* embryos at the 1~4 cell stage at desired concentrations using a pressure-controlled microinjector (WARNER INSTRUMENTS PLI-100A PICO-LITER Injector) and a NARISHIGE micro-manipulator. After injection, the embryos were cultured in E3 embryo medium without methylene blue in the dark at 28.5°C.

### Photo-activation with 3D printed light box

Red light exposure of 6 hpf, 1 dpf and 2 dpf were performed by using a 3D printed LED light box (Figure S1 and Supplementary information). The box was designed using AutoDesk^®^ 123D^®^ Design software and converted to a printable file with scaffolds by Formlabs Preform print preparation software. The converted file was 3D printed by a Formlabs Form 2 printer with black photopolymer resin (FLGPBK03; Somerville, MA), and then trimmed and polished by sand papers. Ventilation windows were includes in the box design to allow air exchange while activation. The red light LEDs (LED660N-03; Roithner LaserTechnik GmbH, Vienna, Austria) and 18 Ω metal film resistors (CCF0718R0GKE36; Vishay, Malvern, PA) were soldered together and connected with a potentiometer (RV24BF-10-15R1-A1K-LA; Alpha, Taiwan) with a 0.87”D Black aluminum knob (405-4764; Eagle Plastic Devices) for adjusting the light intensity (0.98 µW/mm^2^ to 64 µW/mm^2^). A wall mount AC adaptor (WSU135-0620; Triad Magnetics, Perris, CA) is connected to the circuit through a DC power connector (163-4020; Kobiconn). The detailed circuit is shown in Figure. S1A. The light power was measured using an X-Cite^®^ Optical Power Measurement System (Excelitas Technologies Corp.).

### Statistical analysis

All the statistics were calculated by Prism 7 or 8 (GraphPad Software Inc.). An ordinary one way ANOVA was performed and used Dunnett’s multiple comparison test to determine difference between groups.

## ACKNOWLEDGEMENTS

We thank Dr. David Hawk from the Proteomics and Metabolomics Core of MD Anderson Cancer Center for helping the HPLC purification of Phycocyanobilin. We also thank Dr. Adriana Paulucci in the Microscope core of the Genetics Department at the MD Anderson Cancer Center. This work was supported by National Institutes of Health (NIH) grant OD19764 and the Ben F. Love Endowment to R.R.B.; by the NIH training grant, HL007676, to J.C.C; and by the Cancer Prevention Institute of Texas, RR140077, and National Institutes of General Medical Sciences, GM124043, to G.T.E.

## Supplementary information

**Table S1.**
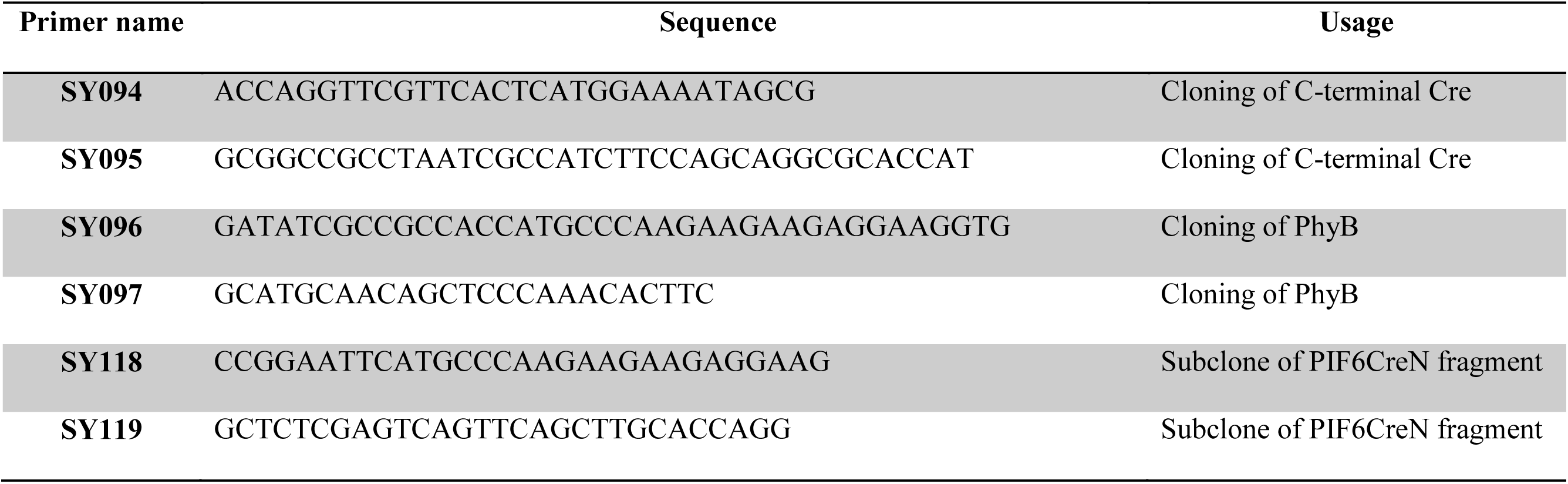
Primers.

**Table S2.** Plasmids.

| Addgene # | Name | Application |
| --- | --- | --- |
| 131780 | pCS2+ PIF6CreN | For *in vitro* transcription of PIF6APBCreN |
| 131781 | pCS2+ PhyBΔCreC | For *in vitro* transcription of PhyBΔCreC |
| 131782 | pME-CreLite | Middle entry clone for Tol2 kit |
| 131783 | pTol2-CreLite | Tol2 transgenesis of CreLite |
| 131785 | pAAV-CreLite | For AAV virus transfection into mammalian tissues |
| 131786 | pLenti-CreLite | For lentivirus transfection in cell culture system |
| 131787 | pTol2-PcyA-IRES-HO1 | Tol2 transgenesis of PCB biosynthesis enzymes |

## Sequences

### 230 bp fragment

AGCCGGAGTCAGCCATGGGAAACTGCGGAAATGGATGCGATTCACTCGCTCCAGCTTATTCTGAGAGACTCTTTTAAAGAATCTGGTGGTTCTGCCGGTGGCTCCGGTTCTGGCTCCAGCGGTGGCAGCTCTGGTGCGTCCGGCACGGGTACTGCGGGTGGCACTGGCAGCGGTTCCGGTACTGGCTCTGGCAACCGGAAATGGTTTCCCGCAGAACCTGAAGATGT TCG

### PhyBΔCreC

ATGCCCAAGAAGAAGAGGAAGGTGGTTTCCGGAGTCGGGGGTAGTGGCGGTGGCCGTGGCGGTGGCCGTGGCGGAGAAGAAGAACCGTCGTCAAGTCACACTCCTAATAACCGAAGAGGAGGAGAACAAGCTCAATCGTCGGGAACGAAATCTCTCAGACCAAGAAGCAACACTGAATCAATGAGCAAAGCAATTCAACAGTACACCGTCGACGCAAGACTCCACGCCGTTTTCGAACAATCCGGCGAATCAGGGAAATCATTCGACTACTCACAATCACTCAAAACGACGACGTACGGTTCCTCTGTACCTGAGCAACAGATCACAGCTTATCTCTCTCGAATCCAGCGAGGTGGTTACATTCAGCCTTTCGGATGTATGATCGCCGTCGATGAATCCAGTTTCCGGATCATCGGTTACAGTGAAAACGCCAGAGAAATGTTAGGGATTATGCCTCAATCTGTTCCTACTCTTGAGAAACCTGAGATTCTAGCTATGGGAACTGATGTGAGATCTTTGTTCACTTCTTCGAGCTCGATTCTACTCGAGCGTGCTTTCGTTGCTCGAGAGATTACCTTGTTAAATCCGGTTTGGATCCATTCCAAGAATACTGGTAAACCGTTTTACGCCATTCTTCATAGGATTGATGTTGGTGTTGTTATTGATTTAGAGCCAGCTAGAACTGAAGATCCTGCGCTTTCTATTGCTGGTGCTGTTCAATCGCAGAAACTCGCGGTTCGTGCGATTTCTCAGTTACAGGCTCTTCCTGGTGGAGATATTAAGCTTTTGTGTGACACTGTCGTGGAAAGTGTGAGGGACTTGACTGGTTATGATCGTGTTATGGTTTATAAGTTTCATGAAGATGAGCATGGAGAAGTTGTAGCTGAGAGTAAACGAGATGATTTAGAGCCTTATATTGGACTGCATTATCCTGCTACTGATATTCCTCAAGCGTCAAGGTTCTTGTTTAAGCAGAACCGTGTCCGAATGATAGTAGATTGCAATGCCACACCTGTTCTTGTGGTCCAGGACGATAGGCTAACTCAGTCTATGTGCTTGGTTGGTTCTACTCTTAGGGCTCCTCATGGTTGTCACTCTCAGTATATGGCTAACATGGGATCTATTGCGTCTTTAGCAATGGCGGTTATAATCAATGGAAATGAAGATGATGGGAGCAATGTAGCTAGTGGAAGAAGCTCGATGAGGCTTTGGGGTTTGGTTGTTTGCCATCACACTTCTTCTCGCTGCATACCGTTTCCGCTAAGGTATGCTTGTGAGTTTTTGATGCAGGCTTTCGGTTTACAGTTAAACATGGAATTGCAGTTAGCTTTGCAAATGTCAGAGAAACGCGTTTTGAGAACGCAGACACTGTTATGTGATATGCTTCTGCGTGACTCGCCTGCTGGAATTGTTACACAGAGTCCCAGTATCATGGACTTAGTGAAATGTGACGGTGCAGCATTTCTTTACCACGGGAAGTATTACCCGTTGGGTGTTGCTCCTAGTGAAGTTCAGATAAAAGATGTTGTGGAGTGGTTGCTTGCGAATCATGCGGATTCAACCGGATTAAGCACTGATAGTTTAGGCGATGCGGGGTATCCCGGTGCAGCTGCGTTAGGGGATGCTGTGTGCGGTATGGCAGTTGCATATATCACAAAAAGAGACTTTCTTTTTTGGTTTCGATCTCACACTGCGAAAGAAATCAAATGGGGAGGCGCTAAGCATCATCCGGAGGATAAAGATGATGGGCAACGAATGCATCCTCGTTCGTCCTTTCAGGCTTTTCTTGAAGTTGTTAAGAGCCGGAGTCAGCCATGGGAAACTGCGGAAATGGATGCGATTCACTCGCTCCAGCTTATTCTGAGAGACTCTTTTAAAGAATCTGGTGGTTCTGCCGGTGGCTCCGGTTCTGGCTCCAGCGGTGGCAGCTCTGGTGCGTCCGGCACGGGTACTGCGGGTGGCACTGGCAGCGGTTCCGGTACTGGCTCTGGCAACCGGAAATGGTTTCCCGCAGAACCTGAAGATGTTCGCGATTATCTTCTATATCTTCAGGCGCGCGGTCTGGCAGTAAAAACTATCCAGCAACATTTGGGCCAGCTAAACATGCTTCATCGTCGGTCCGGGCTGCCACGACCAAGTGACAGCAATGCTGTTTCACTGGTTATGCGGCATAGTGAAACAGGGGCAATGGTGCGCCTGCTGGAAGATGGCGATTAG

NLS

PhyBΔ (1-621 a.a.)

GSAT linker (36 a.a.)

CreC (60 – 343 a.a.)

### PIF6CreN

ATGCCCAAGAAGAAGAGGAAGGTGATGTTCTTACCAACCGATTATTGTTGCAGGTTAAGCGATCAAGAGTATATGGAGCTTGTGTTTGAGAATGGCCAGATTCTTGCAAAGGGCCAAAGATCCAACGTTTCTCTGCATAATCAACGTACCAAATCGATCATGGATTTGTATGAGGCAGAGTATAACGAGGATTTCATGAAGAGTATCATCCATGGTGGTGGTGGTGCCATCACAAATCTCGGGGACACGCAGGTTGTTCCACAAAGTCATGTTGCTGCTGCCCATGAAACAAACATGTTGGAAAGCAATAAACATGTTGACGGTGGTTCTGCCGGTGGCTCCGGTTCTGGCTCCAGCGGTGGCAGCTCTGGTGCGTCCGGCACGGGTACTGCGGGTGGCACTGGCAGCGGTTCCGGTACTGGCTCTGGCCTGACTGTGCACCAAAACCTGCCTGCCCTCCCTGTGGATGCCACCTCTGATGAAGTCAGGAAGAACCTGATGGACATGTTCAGGGACAGGCAGGCCTTCTCTGAACACACCTGGAAGATGCTCCTGTCTGTGTGCAGATCCTGGGCTGCCTGGTGCAAGCTGAACTGA

NLS

PIF6APB (1 – 100 a.a.)

GSAT linker (36 a.a.)

CreN (19 – 59 a.a.)

### PhyBΔCreC-P2A-PIF6CreN

ATGCCCAAGAAGAAGAGGAAGGTGGTTTCCGGAGTCGGGGGTAGTGGCGGTGGCCGTGGCGGTGGCCGTGGCGGAGAAGAAGAACCGTCGTCAAGTCACACTCCTAATAACCGAAGAGGAGGAGAACAAGCTCAATCGTCGGGAACGAAATCTCTCAGACCAAGAAGCAACACTGAATCAATGAGCAAAGCAATTCAACAGTACACCGTCGACGCAAGACTCCACGCCGTTTTCGAACAATCCGGCGAATCAGGGAAATCATTCGACTACTCACAATCACTCAAAACGACGACGTACGGTTCCTCTGTACCTGAGCAACAGATCACAGCTTATCTCTCTCGAATCCAGCGAGGTGGTTACATTCAGCCTTTCGGATGTATGATCGCCGTCGATGAATCCAGTTTCCGGATCATCGGTTACAGTGAAAACGCCAGAGAAATGTTAGGGATTATGCCTCAATCTGTTCCTACTCTTGAGAAACCTGAGATTCTAGCTATGGGAACTGATGTGAGATCTTTGTTCACTTCTTCGAGCTCGATTCTACTCGAGCGTGCTTTCGTTGCTCGAGAGATTACCTTGTTAAATCCGGTTTGGATCCATTCCAAGAATACTGGTAAACCGTTTTACGCCATTCTTCATAGGATTGATGTTGGTGTTGTTATTGATTTAGAGCCAGCTAGAACTGAAGATCCTGCGCTTTCTATTGCTGGTGCTGTTCAATCGCAGAAACTCGCGGTTCGTGCGATTTCTCAGTTACAGGCTCTTCCTGGTGGAGATATTAAGCTTTTGTGTGACACTGTCGTGGAAAGTGTGAGGGACTTGACTGGTTATGATCGTGTTATGGTTTATAAGTTTCATGAAGATGAGCATGGAGAAGTTGTAGCTGAGAGTAAACGAGATGATTTAGAGCCTTATATTGGACTGCATTATCCTGCTACTGATATTCCTCAAGCGTCAAGGTTCTTGTTTAAGCAGAACCGTGTCCGAATGATAGTAGATTGCAATGCCACACCTGTTCTTGTGGTCCAGGACGATAGGCTAACTCAGTCTATGTGCTTGGTTGGTTCTACTCTTAGGGCTCCTCATGGTTGTCACTCTCAGTATATGGCTAACATGGGATCTATTGCGTCTTTAGCAATGGCGGTTATAATCAATGGAAATGAAGATGATGGGAGCAATGTAGCTAGTGGAAGAAGCTCGATGAGGCTTTGGGGTTTGGTTGTTTGCCATCACACTTCTTCTCGCTGCATACCGTTTCCGCTAAGGTATGCTTGTGAGTTTTTGATGCAGGCTTTCGGTTTACAGTTAAACATGGAATTGCAGTTAGCTTTGCAAATGTCAGAGAAACGCGTTTTGAGAACGCAGACACTGTTATGTGATATGCTTCTGCGTGACTCGCCTGCTGGAATTGTTACACAGAGTCCCAGTATCATGGACTTAGTGAAATGTGACGGTGCAGCATTTCTTTACCACGGGAAGTATTACCCGTTGGGTGTTGCTCCTAGTGAAGTTCAGATAAAAGATGTTGTGGAGTGGTTGCTTGCGAATCATGCGGATTCAACCGGATTAAGCACTGATAGTTTAGGCGATGCGGGGTATCCCGGTGCAGCTGCGTTAGGGGATGCTGTGTGCGGTATGGCAGTTGCATATATCACAAAAAGAGACTTTCTTTTTTGGTTTCGATCTCACACTGCGAAAGAAATCAAATGGGGAGGCGCTAAGCATCATCCGGAGGATAAAGATGATGGGCAACGAATGCATCCTCGTTCGTCCTTTCAGGCTTTTCTTGAAGTTGTTAAGAGCCGGAGTCAGCCATGGGAAACTGCGGAAATGGATGCGATTCACTCGCTCCAGCTTATTCTGAGAGACTCTTTTAAAGAATCTGGTGGTTCTGCCGGTGGCTCCGGTTCTGGCTCCAGCGGTGGCAGCTCTGGTGCGTCCGGCACGGGTACTGCGGGTGGCACTGGCAGCGGTTCCGGTACTGGCTCTGGCAACCGGAAATGGTTTCCCGCAGAACCTGAAGATGTTCGCGATTATCTTCTATATCTTCAGGCGCGCGGTCTGGCAGTAAAAACTATCCAGCAACATTTGGGCCAGCTAAACATGCTTCATCGTCGGTCCGGGCTGCCACGACCAAGTGACAGCAATGCTGTTTCACTGGTTATGCGGCGGATCCGAAAAGAAAACGTTGATGCCGGTGAACGTGCAAAACAGGCTCTAGCGTTCGAACGCACTGATTTCGACCAGGTTCGTTCACTCATGGAAAATAGCGATCGCTGCCAGGATATACGTAATCTGGCATTTCTGGGGATTGCTTATAACACCCTGTTACGTATAGCCGAAATTGCCAGGATCAGGGTTAAAGATATCTCACGTACTGACGGTGGGAGAATGTTAATCCATATTGGCAGAACGAAAACGCTGGTTAGCACCGCAGGTGTAGAGAAGGCACTTAGCCTGGGGGTAACTAAACTGGTCGAGCGATGGATTTCCGTCTCTGGTGTAGCTGATGATCCGAATAACTACCTGTTTTGCCGGGTCAGAAAAAATGGTGTTGCCGCGCCATCTGCCACCAGCCAGCTATCAACTCGCGCCCTGGAAGGGATTTTTGAAGCAACTCATCGATTGATTTACGGCGCTAAGGATGACTCTGGTCAGAGATACCTGGCCTGGTCTGGACACAGTGCCCGTGTCGGAGCCGCGCGAGATATGGCCCGCGCTGGAGTTTCAATACCGGAGATCATGCAAGCTGGTGGCTGGACCAATGTAAATATTGTCATGAACTATATCCGTAACCTGGATAGTGAAACAGGGGCAATGGTGCGCCTGCTGGAAGATGGCGATGGAAGCGGAGCTACTAACTTCAGCCTGCTGAAGCAGGCTGGAGACGTGGAGGAGAACCCTGGACCTATGCCCAAGAAGAAGAGGAAGGTGATGTTCTTACCAACCGATTATTGTTGCAGGTTAAGCGATCAAGAGTATATGGAGCTTGTGTTTGAGAATGGCCAGATTCTTGCAAAGGGCCAAAGATCCAACGTTTCTCTGCATAATCAACGTACCAAATCGATCATGGATTTGTATGAGGCAGAGTATAACGAGGATTTCATGAAGAGTATCATCCATGGTGGTGGTGGTGCCATCACAAATCTCGGGGACACGCAGGTTGTTCCACAAAGTCATGTTGCTGCTGCCCATGAAACAAACATGTTGGAAAGCAATAAACATGTTGACGGTGGTTCTGCCGGTGGCTCCGGTTCTGGCTCCAGCGGTGGCAGCTCTGGTGCGTCCGGCACGGGTACTGCGGGTGGCACTGGCAGCGGTTCCGGTACTGGCTCTGGCCTGACTGTGCACCAAAACCTGCCTGCCCTCCCTGTGGATGCCACCTCTGATGAAGTCAGGAAGAACCTGATGGACATGTTCAGGGACAGGCAGGCCTTCTCTGAACACACCTGGAAGATGCTCCTGTCTGTGTGCAGATCCTGGGCTGCCTGGTGCAAGCTGAACTGA

PhyBΔCreC (60-343 a.a.)

GSG-P2A (Kim et al., 2011)

PIF6APBCreN

## Supplementary Methods

### Phycocyanobilin (PCB) handling

PCB has to be handled with care. Previous study have shown that PCB is prone to oxidation and will change its color from dark blue to brown in cell culture media after 24 hours (Müller et al., 2014). In addition, methylene blue, a common antifungal additive in zebrafish medium, also a common indicator for redox reaction, may be a potential oxidant to eliminate the activity of PCB. An addition of an anti-oxidant or reducing agents like β-mercapto-ethanol may help to limit the oxidation of PCB in culture. Notably, Buckley et al. (Buckley et al., 2016) has shown that the HPLC-purified PCB from cyanobacterium Spirulina extraction is toxic at higher dose of PCB (10 µM). However, the toxicity effect has not been shown when we use chemically synthesized PCB at 14 µM. It is possible that the toxicity arises from components in cyanobacteria instead of PCB itself.

**Supplementary figure 1.**
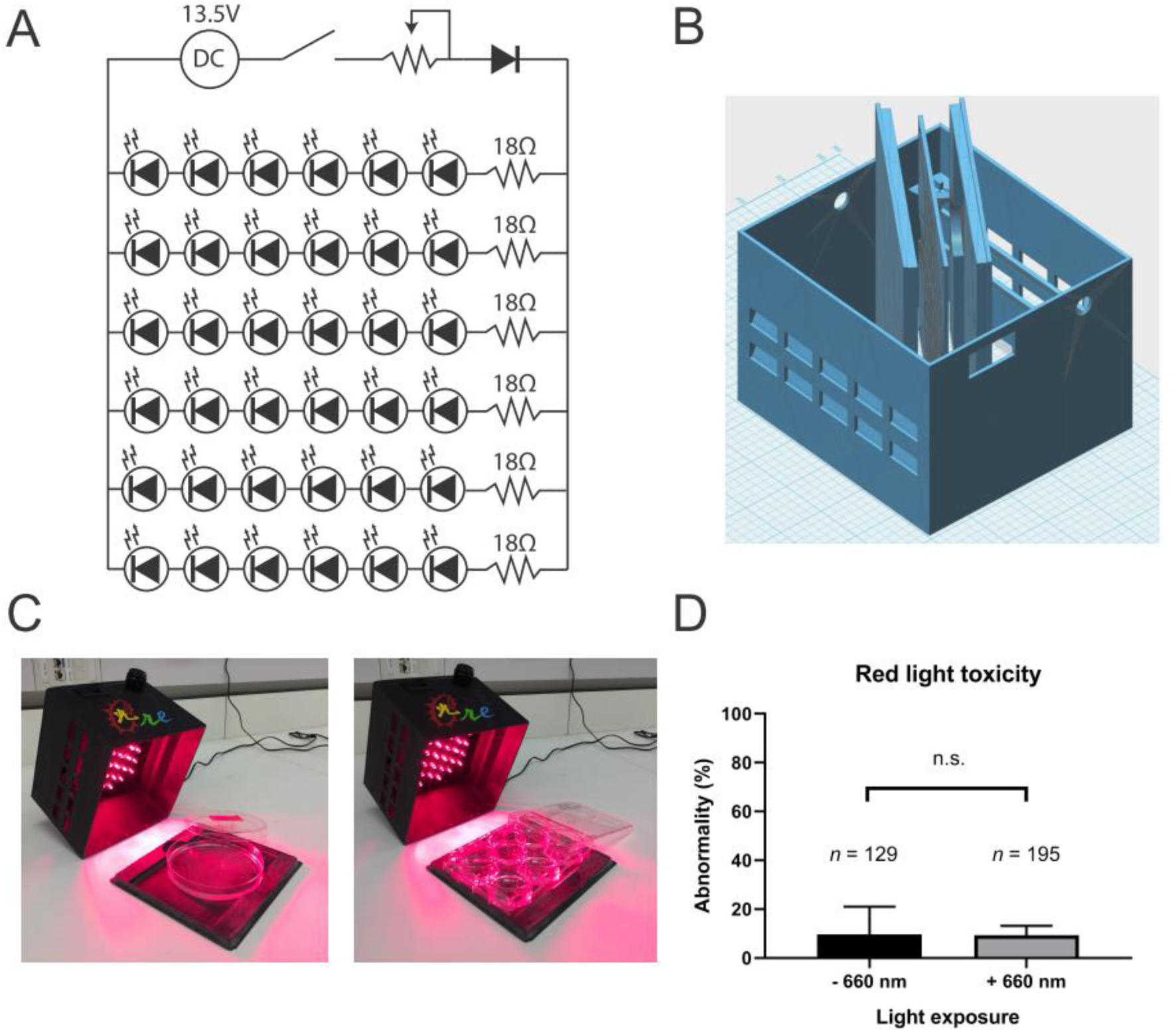
Illumination and red-light box design. The red LED light box was 3D printed. High power LEDs, resistors, a rocker switch, and a potentiometer were soldered together (A) on a 3D-printed resin circuit board. The illumination box has slits to ensure air-flow into the box, but also blocks the light path into the bottom of the box (B). The bottom of the illumination box can hold either a 100 mm petri dish or a 6-well plate, allowing large area activation (C). Mean number of embryos with abnormality with or without red light (660 nm) exposure (D). Data are from three independent experiments and error bars represent *SD*; paired, two-tailed *t*-test. *n* = total number of embryos in each group.

